# Maintenance of Bound or Independent Features in Visual Working Memory is Task-dependent

**DOI:** 10.1101/2024.10.22.619709

**Authors:** Ruoyi Cao, Leon Y. Deouell

## Abstract

Conflicting findings exist regarding whether features of an object are stored separately or bound together in visual working memory. This controversy is based on an implicit assumption about a default, or fixed, mode of working memory storage. In contrast, here we asked whether the anticipated task might determine the format in which information is maintained in working memory, consistent with its task-oriented function. To test this flexible maintenance hypothesis, we recorded EEG while subjects performed a delayed (Yes/No) recognition task with different requirements and loads. Across three experiments, we compared event-related potentials (ERPs) in conditions with and without the necessity of maintaining conjunctions between features, while controlling for differences in visual stimulation. In Experiment 1 (N=24), involving color-location conjunctions, we identified a delay-period effect characterized by a positive potential shift in central-parietal channels when conjunction was not required by the task. This pattern, distinct from the effect caused by an increased working memory load, was confirmed in Experiment 2 (N=23) with an independent group of subjects using a similar paradigm, while also controlling for the physical appearance of probes. Finally, the observation of color and location conjunction in Experiments 1 and 2 was extended to Color and Orientation conjunction in Experiment 3 (N=22). Collectively, these three experiments provided reliable evidence demonstrating that the maintenance of feature conjunctions in working memory, whether spatial (location) or non-spatial (non-location), depends on the task goal.

**Highlights:**

1. Task demands determine whether objects are represented as separate features or as conjoined items (maintenance format) in visual working memory.
2. EEG recordings show differential activity despite similar stimulation, reflecting task demands.
3. The effect of the task is evident in the early stages of working memory processing.
4. Flexibility in maintenance format was observed for both color-location and color-orientation conjunctions.
5. The findings challenge the notion of fixed maintenance format during visual working memory.

## 1. Introduction

Visual input is composed of various visual features, such as color, location, shape, orientation, and movement. To interact with the world efficiently, we need to not only correctly integrate different features into an object but also establish an internal representation of bound objects that remains accessible after the visual stimuli disappear, that is, visual working memory (VWM). The question of whether features are stored bound or separately in working memory has garnered significant attention in both psychology and neuroscience (Treisman & Gelade, 1980; Treisman, 1998; Schneegans & Bays, 2019; Treccani, 2018).

A typical, if indirect, way to investigate this question is by correlating performance in delayed recognition or match-to-sample tasks with the number of items to be remembered. If features are separately stored in working memory, then performance should be limited by the number of features rather than the number of objects to be maintained. Otherwise, if bound objects are stored, performance should be limited by the number of objects regardless of the number of features. Following such reasoning, Luck and Vogel (1997) proposed that features are conjoined as items in VWM, as they observed that increasing the number of features within objects did not impair performance in change-detection tasks. Moreover, invariance to the number of features was observed even when one object consisted of multiple values from the same feature dimension (like multiple colors). Such a perspective, referred to as the ‘strong object’ view, was also supported by a study showing no difference in delay EEG activity between the memorization of single- or multi-color objects (Luria & Vogel, 2011).

However, other studies came to different conclusions. For example, Wheeler and Treisman (2002) observed that while features from different dimensions could be stored in parallel without a cost for working memory performance, adding features from the same feature dimension limited memory performance. They proposed the ‘multiple-resources’ view, stating that VWM maintains features from different feature dimensions in parallel, while features from the same feature dimension compete for storage space. The multiple-resources view was later supported by other studies using either similar (Delvenne & Bruyer, 2004) or different paradigms (Wang, 2017, Xu, 2002). For example, Wang et al (2017) found that for objects with color and orientation, the memory performance for color decreased as more colors had to be remembered, but the memory for orientation was not affected, and vice versa for orientation

Yet another model proposes that VWM performance is limited by both the number of objects and the number of features. Olson and Jiang (2002) found that when the number of objects was held constant, performance was superior in a single-feature condition compared to a multiple-feature condition, indicating greater difficulty in storing two features than one feature of an object. Conversely, when the number of features was held constant, performance was better when features conjoined to form objects than when they were presented as isolated features. The importance of both the number of features and the number of items inspired the ‘weak-objects view’ of working memory (Alvarez & Cavanagh, 2004; Hardman & Cowan, 2015). Imaging studies have also supported both the maintenance of separate features and bound objects. For instance, both non-human primate studies (Baizer, Ungerleider, & Desimone, 1991; Mishkin, Ungerleider, & Macko, 1983) and human studies (Courtney, Ungerleider, Keil, & Haxby, 1996; Smith et al., 1995) found that VWM for spatial location and item identities activated different regions of the brain. Other studies however found no evidence for distinct representations of spatial and non-spatial features (D’Esposito et al., 1998; Kravitz, Kriegeskorte, & Baker, 2010; Kravitz, Saleem, Baker, & Mishkin, 2011).

Debates about whether features are stored separately or bound are based on an implicit assumption that there is a default or fixed mode of VWM storage. However, the existence of such a universal mode cannot be presumed, given the remarkable flexibility in prioritizing information in WM according to the task goal (Freedman et al., 2001; Duncan, 2001; McCants et al., 2019). For example, when non-human primates viewed the same to-be-remembered stimuli but were trained to expect different kinds of memory probes, delay activity in the prefrontal cortex showed different patterns (Rao, Rainer, & Miller, 1997). In an fMRI study in humans, participants were required to remember a face and a scene (Nobre, 2007). During the delay period, a cue was presented to inform which is to be tested later. Increased activity was observed in areas involved in the face (fusiform gyrus) or scene (Parahippocampal gyrus) processing, according to the cue. Indeed, VWM is not simply a representational state of visual input during a delay period but is better conceived as a functional state bridging previous contexts and sensations to anticipated actions and outcomes (Myers, Stokes, & Nobre, 2017). Consequently, the anticipated memory probes (questions) used in a given experiment might determine the format in which objects will be maintained in VWM and the involved cognitive resources.

To test the Flexible Maintenance hypothesis, proposing that the brain stores separate features or bound objects based on the task goal, EEG recordings were obtained while subjects engaged in a delayed (Yes/No) recognition task, both with and without the requirement of maintaining binding between features. If there is a fixed mode for the system to store visual stimuli regardless of task requirement, we expected no systematic differences between these two conditions during the delay period, as the visual input to be maintained was identical. In contrast, if information maintained in VWM is adaptive to task goals, we expected to find such differences. Three experiments with similar paradigms were conducted using a delay recognition task in which participants viewed a small array of objects and following a delay had to decide if a probe was similar to one of the objects in the array or not. We compared EEG patterns during conditions with and without the need to maintain information about the conjunction between two features. In Experiment 1, we investigated the impact of task requirements to maintain conjunctions between color and location, following identical visual input during the encoding and delay period. In Experiment 2, we replicated the findings of Experiment 1 while also controlling the physical appearance of the probe. In Experiment 3, we extended this experiment into binding between colors and orientations.

## 2 Experiment 1

### 2.1 Methods

#### 2.1.1 Participants

Thirty-five healthy volunteers from the Hebrew University of Jerusalem participated in the study. Subjects were paid (40NIS/h, ∼$12) or given course credits for participation. All subjects had reportedly normal or corrected-to-normal sight and no psychiatric or neurological history. One subject did not finish the experiment and nine were excluded from analysis due to noisy recordings according to the criteria described below. The remaining twenty-four subjects consisted of 12 males and 12 females (19–31 years old). The experiment was approved by the ethics committee of the Hebrew University of Jerusalem, and informed consent was obtained for each subject after the experimental procedures were explained.

#### 2.1.2 Stimuli and Apparatus

Subjects sat in a dimly lit room. The stimuli were presented with Psychotoolbox-3 (http://psychtoolbox.org/) implemented in Matlab 2018 (The MathWorks, Inc., Natick, Massachusetts) on a ViewSonic G75f CRT (1024×768) monitor with a 100-Hz refresh rate. They appeared on a grey background at the center of the computer screen located 100 cm away from the subjects’ eyes.

Subjects performed a delayed (Yes/No) recognition test in which a memory array was followed by a probe with a delay of 1-3 seconds. Each subject performed 3 blocks of trials from 3 conditions (Figure 1a): two-item-feature (F2), two-item-binding (B2), and four-item-binding (B4). The memory array consisted of 2 colored items in the F2 and B2 conditions and 4 items in the B4 conditions. All items had an identical irregular shape, which subtended a visual angle of 2.1×2.1°. The color of each item was randomly selected out of six highly distinguished colors, including red (RGB: 255,0, 0), green (0, 255, 0), yellow (255,255, 0), blue (0, 0, 255), violet (255,255,0) and white (0,0,0), without repetition (i.e. within a given array, each item had a unique color). The locations of the items were randomly selected out of eight potential locations evenly distributed on an invisible circle with a diameter of 7.3°centered on the fixation cross. Probes consisted of a single item, which could be a feature probe or a binding probe, of different match states, as described in Table 1. Binding probes were color shapes in the periphery. In contrast, feature probes consisted of a black dot in the periphery, or a color shape in the center (see examples in Figure 1a).

**Figure 1.**
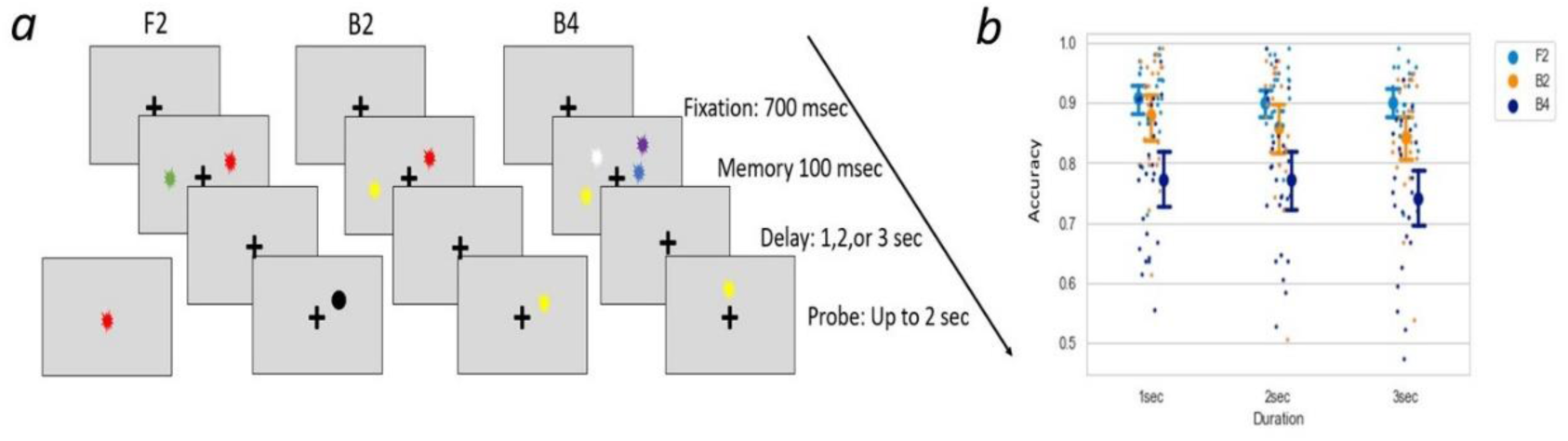
(a) Illustration of an example trial in the three conditions: Left – Two-Item feature condition (F2) with a matched probe; middle – Two-Item binding condition (B2) with a mis-conjunction probe; right – Four-Item binding condition (B4) with a new-feature probe. (b) Percentage of correct responses for each condition following 1, 2, or 3-second delays between the learning and test phases. Here and in subsequent figures, small dots represent individual subject means, while large dots and error bars depict across-subject means and 95% confidence intervals.

**Table 1:** Description of probes used in Experiment 1.

| Probe type | Match State | Description | Correct Response | Trial Number |
| --- | --- | --- | --- | --- |
| Binding probe | Matched | A probe with the same color and location as one of the items in the memory array. | Matched | 144 |
|  | New-location | A probe with the same color but in a different location | Non-matched | 24 |
|  | New-color | A probe in the same location but with a different color | Non-matched | 24 |
|  | Mis-conjunction | A probe with both color and location presented in the memory array but switched between items | Non-matched | 96 |
|  | Neither-matched | A probe with neither color nor location presented in the memory array | Non-matched | 24 |
| Feature Probe | Color-matched | A probe in the center of the screen with a color presented in the memory array | Matched | 72 |
|  | New-color | A probe in the center of the screen with a color not presented in the memory array | Non-matched | 72 |
|  | Location-matched | A probe presented as a black circle at the location of one of the items in the memory array. | Matched | 72 |
|  | New-location | A probe presented as a black circle at a location not matching any of the items in the memory array | Non-matched | 72 |

#### 2.1.3 Experimental procedure

Each trial started with a 700 msec fixation cross presented at the center of the screen (Figure 1a). Next, the memory array appeared for 100 msec. The memory array was followed by a delay period of either 1,2 or 3 seconds of blank screen, in equal proportions. Trials with the 3 delay durations were randomly mixed within a block. Following the delay period, a probe item was presented, and subjects had to press a key to indicate if the probe item was a “Matched” or a “Non-matched” item. The probe disappeared when the response was made (with a limit of 2 seconds), followed by the initial fixation of the subsequent trial. In the F2 condition, subjects did not have to consider the conjunction of color and location, and only decide whether the color or the location (depending on the probe type) were in the memory array or were novel. In the B2 and B4 conditions, subjects had to also consider the conjunction of color and location and respond to mis-conjunctions as “Non-matched”.

There were 3 conditions in the experiment, each condition consisting of 288 trials. In the B2 and B4 conditions, the delay period was followed by Match probes in 144 trials, and non-match probes in the remaining trials: *New-color probes* in 24 trials, *new- location probes* in 24 trials, and *Mis-conjunction* probes in 96 trials. In the F2 condition, the delay period was followed by 72 trials of each type (*Color-matched, New-color, Location-matched, New-location).* Each condition began with 64 practice trials followed by 2 consecutive blocks of that condition, resulting in 6 blocks in total (288 trials for each condition). Each block took about 15 mins, and subjects were instructed to take a break between blocks. The order of the three conditions was counterbalanced across participants.

As the memory arrays were identical in B2 and F2 conditions, containing 2 items each with 2 relevant features, the contrast between B2 and F2 was intended to reveal the impact of individual feature values and their conjunctions, denoted as the Task effect. The comparison between B4 and B2 was conducted to unveil a Load effect, specifically focusing on the retention of 4 items versus 2 items.

### 2.2 Analysis

#### 2.2.1 Behavior

Statistical analysis was conducted with JASP (Version 0.17.1; https://jasp-stats.org/) and figures were made by the Python graphic library Seaborn (https://www.python-graph-gallery.com/seaborn/). Response accuracy of subjects was entered into a 3× 3 Delay duration (1 vs 2 vs 3 sec delay) by Condition (F2 vs B2 vs B4) repeated-measures two-way ANOVA. Degrees of freedom were adjusted for violations of the assumption of sphericity with the Greenhouse–Geisser (GG) correction when necessary, and the uncorrected degrees of freedom were reported together with the GG epsilon.

#### 2.2.2 EEG recording and data analysis

EEG data were recorded using an Active 2 system (Biosemi, the Netherlands) from 64 active electrodes spread out across the scalp according to the extended 10–20 system with the addition of two mastoid electrodes and a nose electrode (https://www.biosemi.com/pics/cap_64_layout_medium.jpg). Horizontal electro- oculogram (EOG) was recorded from electrodes placed at the outer canthi of both eyes. Vertical EOG was recorded from electrodes placed above and below the left eye. The EEG was continuously sampled at 1024 Hz with an anti-aliasing low pass filter with a cutoff of 1/5 the sampling rate, and stored for off-line analysis. The data was referenced online to the Common Mode Signal (CMS) electrode which was placed in the space between POz, PO3, Pz, and P1.

Data preprocessing and analysis were performed using the FieldTrip toolbox (Oostenveld et al., 2011; http://fieldtriptoolbox.org) implemented in MATLAB R2018 (MathWorks, Natick, MA, USA). Preprocessing was applied to continuous data. During preprocessing, EEG and EOG signals were first filtered with a 4th-order Butterworth zero-phase (forward and reverse filter) bandpass filter of 0.1–180 Hz and then referenced to the nose channel. Extremely noisy or silent channels, which contributed more than 20% of all artifacts (Defined as more than 100μV absolute difference between samples within segments of 100 msec or absolute amplitude > 100μV) were deleted. Subjects who lost either more than 2 adjacent channels or more than 3 channels were excluded. Next, data were re-referenced to an average of all remaining EEG electrodes. Ocular and muscular artifacts were removed from the EEG signal using the ICA method by manual selection of artifact components based on correlation with the EOG channels, power spectrum, and component scalp topographies (Jung et al., 2000).

After ocular and muscle artifacts were removed by ICA, automatic artifact rejection was applied (http://www.fieldtriptoolbox.org/tutorial/automatic_artifact_rejection/) with the following criteria. Time points larger than 12 standard deviations from the mean of the corresponding channel were marked, together with 200 msec before and after, so that the (subthreshold) beginning and end of an artifactual event will be likely accounted for. A visual inspection of the data followed to detect rare artifacts that were missed by the automatic procedure. Finally, previously deleted channels were recreated by mean interpolation of the neighboring electrodes (See Supplementary Table S1 for the deleted channels for each subject in Experiment 1). Data were then down-sampled to 512 Hz, filtered with a Butterworth zero-phase lowpass filter at a cutoff at 20 Hz, and parsed into 1800 msec segments starting 500 msec before the memory array onset. Trials overlapping time points marked as artifact in the previous steps were excluded from analysis.

We initially analyzed the first 11 subjects, and then replicated the results on the subsequent group of 13 subjects using the same paradigm and setup (Cao, Pertzov, Gao, Shen, & Deouell, 2021). Since no significant differences were found after a 1-second delay in our previous analysis, we used the first second (shared by all trials) for further analysis. In this paper, we applied a new, computationally efficient implementation of the linear mixed model recently suggested by Visalli et al. (2024) on the entire group of 24 subjects (see next section). This approach allowed us to include single trials, rather than subject averages, thereby enhancing sensitivity. We followed this exploratory analysis with a replication on a new group of subjects (Experiment 2).

#### 2.2.3 Linear-mixed model

Although being a powerful method, the adoption of linear-mixed models in neuroimaging analysis has been limited due to the huge computational demands for mass univariate linear analysis with permutation. In our analysis, we employed a recently published method by Visalli et al (2024) that speeds the computation by approximately a factor of 300. Our data were analyzed with the function that implemented the Mass linear mixed-effects modeling of EEG data with crossed random effects provided by the associated toolbox (ImeEEG: github.com/antovis86/lmeEEG).

The average of 100 msec before the onset of the memory array was defined as a baseline and subtracted from all data points of each segment. To reduce the computational load for this analysis, we down-sampled the data again to 256 Hz and averaged over groups of three consecutive trials, maintaining alignment with the presentation sequence (For example the 1st, 2nd, and 3rd trials were averaged and created the 1st trial in these averaged trial data). All 64 EEG channels were included, and we selected the time points from the beginning of the delay period (the offset of the memory array) until 1000 msec thereafter. The voltage of each channel and each time point was modeled using a separate linear-mixed model to examine the relationship between voltage measurements and experimental conditions, controlling for individual differences and task-irrelevant changes over time.

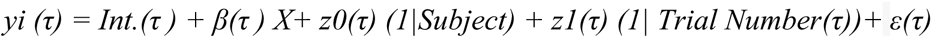

where

- *yi (τ)*: voltage at channel i during time τ;
- *Int. (τ)*: intercept at time τ, capturing the baseline voltage shared across experimental conditions.
- *X*: the experimental condition (B2 vs F2; B2 vs B4).
- *β(τ)*: the coefficient associated with the effect of the condition at time τ.
- *z0(τ) (1|Subject)*: subject-specific variations in baseline voltage at time *τ*.
- *z1(τ) (1| Trial Number(τ)*: task-irrelevant changes across recording time at time τ
- ε(τ): the residual, accounting for unexplained variability.

The experimental condition (B2 vs F2/ B2 vs B4) was considered as the fixed factor, allowing assessment of the effect of interest (Task effect /Load effect) on voltage. Random effects, such as *(1/Subject) and (1Trial Number(τ))*, respectively accounted for individual differences and task-irrelevant changes across recording time. The resulting t statistics map (t-mapsObs in the ImeEEG toolbox), calculated as the squared ratio of the fixed factor estimates to their standard errors, provided insight into the significance of the estimation. The t-statistics were compared with the permutation distribution of the t-statistics obtained by randomly switching the condition labels for 2000 iterations. During the permutation procedure, instead of repeatedly fitting the raw (shuffled) data with the linear mixed model, the "marginal" EEG (mEEG) data— reconstructed by removing the fitted random values (*z0(τ) (1|Subject) + z1(τ) (1|Trial Number)*) from the raw EEG data—was fitted with a multiple linear regression (LM) without the random effect. This approach yields the same results for fixed effects but with significantly faster computation compared to a linear mixed model that includes random effects (Visalli et al., 2024). A p-value is obtained by comparing the t-statistics of the data to the distribution generated by the permutation procedure. The false discovery rate (FDR) was used to correct for family-wise error across time points (Benjamini & Hochberg, 1995).

#### 2.2.4 MVPA

In addition to directly contrasting ERPs between conditions, multivariate pattern analysis (MVPA) was applied. If the experimental condition can be successfully “decoded” from the participant’s patterns of brain activation, we could conclude that information relevant to the experimental manipulation existed in such multidimensional data at that moment.

In addition to training and testing classifiers to distinguish B2 vs F2 condition (Task effect) and B4 vs B2 condition (Load effect ) respectively, we used cross-condition decoding to check whether the “Task effect” revealed by contrast between B2 vs F2 shared the same underlying patterns as Load effect revealed by B2 vs B4 contrast. If classifiers trained and tested across different contrasts performed above chance level, it would suggest that both contrasts might reflect the same effect. On the other hand, if cross-decoding performance hovered around the chance level, even though successful decoding was achieved within contrast conditions, it would support the distinctiveness of the Task and Load effects.

MVPA was performed in MATLAB using the MPVA light toolbox (https://github.com/treder/MVPA-Light). The classification was performed for each participant in a time-resolved manner. Trials for each participant in each condition were first sub-sampled to the minimum number across the three conditions to obtain a balanced set for training and testing. These data were then split into five equal folds. For each time point, a classifier was trained on four folds, and tested on the remaining fold. Each fold was used as a testing set once, and the final accuracies were averaged over the five folds. To increase the signal-to-noise ratio, we randomly sampled groups 5 trials to create super-trials for the training and testing sets separately (i.e. trials from the training set were never mixed with the trials from the test set, to avoid overfitting). After each selection, the trials were returned to the pool. This process was repeated until the number of super-trials equaled the number of trials before averaging. A linear discriminant (LDA) classifier was trained for each time point from 100 msec before the memory array onset to 1000 msec after. To attain decoding performance independent of the decision boundary, the Area Under the receiver operating Curve (AUC) was calculated for each subject. To reveal the generalization of the resulting classifiers over time, a temporal generalization matrix, where classifiers were trained on one time point and tested on all time points, was computed for each subject (King & Dehaene, 2014).

For each decoding analysis, we compared the obtained AUC with chance (AUC = 0.5), using False Detection Rate (FDR) to correct for multiple comparisons at the group level.

### 2.3 Results

#### 2.3.1 Behavioral results

A 3 × 3 Delay duration (1 vs 2 vs 3 sec) × Condition ( 2F vs 2B vs 4B) repeated-measures two-way ANOVA of response accuracy revealed a main effect of Condition, *F(2,46)* = 29.24, *p* < .001, *η2* = 0.56 (Figure 1b). Follow-up pairwise comparison (with Bonferroni correction) showed a significantly lower response accuracy in condition B4 than both condition B2, *Mdiff = - 0.099, SE = .02, t(23) = -5.21, p < .001, Cohen’d =- 1.06*, and condition F2, *Mdiff = - .14, SE = .02, t(23) = -7.46, p < .001, Cohen’d = - 1.52*, whereas no significant differences were found between B2 and F2 condition, *Mdiff = - .04, SE = .02, t(23) = -2.25, p = .09*. The main effect of Delay Duration was also significant, *F(2,46) = 6.07, p = .005, η2 =.21*. Follow-up pairwise comparisons showed a significant decrease of response accuracy only from the 1 sec to the 3 sec delay, *Mdiff* = -0.02, *SE* = .01, *t(23)* = - 3.44, *p* = .004, *Cohen’d* = .70, reflecting some memory decay following extended delays. No interaction between Delay Duration and the Condition was found*, F(4,92)* =1.51, *p* = 0.21. These results indicated that the B4 condition, as expected, was more difficult than the other conditions, and that F2 and B2 conditions did not significantly differ in their overall difficulty.

#### 2.3.2 Linear-Mixed model results

To reveal the Task effect, we compared ERP amplitudes in conditions where subjects were presented with identical memory arrays consisting of two items. One condition required maintaining the conjunction between color and location (B2), while the other did not (F2). We found that the Feature condition elicited significantly more positive voltage responses compared to the Binding condition at central-parietal channels (See Figures 2a and 2c for latencies), and more negative voltage responses compared to the Binding condition at the frontal channels.

**Figure 2:**
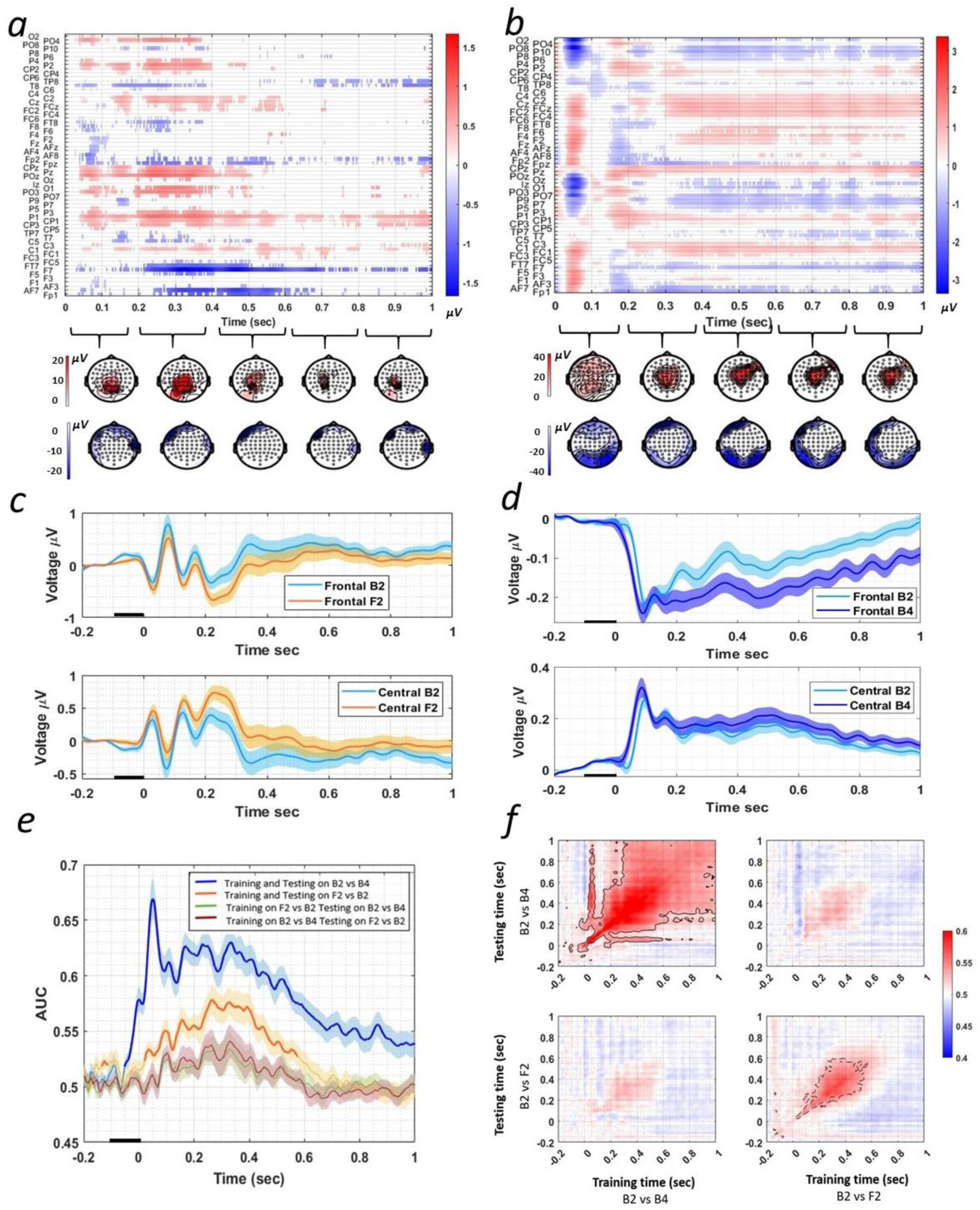
ERP results of Experiment 1. (a) Top Panel: Difference between the F2 and B2 conditions across channels (rows) and time, where significant (p < 0.05, FDR corrected) positive differences are displayed in red and negative differences in blue. Here and in the other panels, Time 0 is the beginning of the delay period. Bottom Panel: Difference topographies between the F2 and B2 conditions, with colors representing the sum of significant voltage differences over each 200 ms interval (p < 0.05, FDR corrected). The top row shows the positive differences, and the bottom row shows the negative differences. (b) Same as (a) for the B4 vs B2 difference. (c) Average event-related potentials (ERPs) triggered by the F2 and B2 conditions, averaged across subjects. The top panel displays ERPs from central channels showing significant positive differences, while the bottom panel shows ERPs from frontal channels showing significant negative differences. In this and subsequent ERP figures, shading around the waveforms indicates the standard error of the mean across subjects. The horizontal black bar indicates the duration of the memory array. (d) Same as (c) for the B4 vs B2 difference. (e) Decoding performance (AUC) in Experiment 1, showing decoding within and across contrasts. Thick line segments indicate intervals with significant above-chance decoding (compared to AUC = 0.5, FDR corrected, p < 0.01). The shaded area represents the standard error across subjects. A black bar on the x-axis represents the presentation of the memory array. (f) Temporal generalization matrices within and across contrasts. Testing time is shown on the vertical axis and training time on the horizontal axis. Significant decoding is indicated by black contours (p < 0.01, FDR corrected). Note that the AUC lines in panel (e) are the diagonals of the matrices in panel (f), respectively.

To reveal the Load effect, we compared the ERP amplitudes between conditions requiring the retention of four bound items (B4) and two bound items (B2). We found that a higher Load elicited more positive voltage at the central channels and more negative voltage surrounding it (See Figures 2b and 2d for latencies).

#### 2.3.3 MPVA results

Above chance decoding of B2 vs F2 was found from 20 msec after the onset of the delay (120 msec after the onset of the memory array) until 560 msec, with the peak accuracy at 265 msec after the memory array offset (Figure 2e; FDR corrected, p<0.01). As the B2 and F2 conditions had similar memory arrays with two items, such above chance decoding reflects the Task effect. Significant decoding performance (AUC) between B2 and B4 was found from 40 msec before the onset of delay (60 msec after the onset of the memory array, *p* < 0.01 FDR corrected) lasting until the end of the delay period (1000 msec after the delay onset) with the peak accuracy at 52 msec. This above-chance decoding reflects the Load effect as well as visual differences between the arrays containing either two or four items. Subsequently, we investigated whether a classifier trained to differentiate between B2 and B4 could decode B2 and F2, and vice versa. Successful cross-decoding would imply similar spatial distributions of EEG responses between the Load effect and the Task effect. However, even though we could decode B2 from F2 and B2 from B4 significantly above chance, we were not able to find a significant above-chance decoding when training and testing on different contrasts (Figure 2e).

One possible cause of the absence of cross-decoding between contrasts for the Load and Task effect is that these effects might exhibit distinct temporal dynamics, where the same spatial distribution manifests in different periods. To explore this hypothesis, classifiers were trained at each time point and subsequently tested on all time points. Figure 2f shows the temporal generalization metrics averaged over subjects. No notable cross-contrast decoding emerged in the temporal generalization matrix as well (top-right and bottom-left panels, Figure 2f). This implies that even when accounting for potential different temporal dynamics, there is no substantial evidence suggesting that the Task effect is the same effect caused by working memory load. We observed also that the two effects have different generalization patterns (top-left and bottom-right panels, Figure 2f)

### 2.4 Interim discussion

With both mass Linear Mixed-Effects Modeling and MVPA, we affirmed that distinct cognitive demands for feature binding into objects yield different activity patterns starting early during the memory delay. As the memory array was identical in F2 and B2 conditions, a difference between the two conditions could not be due to differences in visual information. Additionally, such differences could not be completely accounted for by effects caused by working memory load: performance was similar across the F2 and B2 conditions, and the load effect did not generalize to the task effect. This provides evidence of task-dependent encoding and maintenance processes and argues against the notion, implicitly embedded in previous literature, that a fixed default mode of item representation, either bound or not, is maintained in working memory.

However, while the memory arrays were identical in the F2 and B2 conditions, the probe was different. Even though the differences in ERPs and the decoding results refer to the time before the probe was presented, given that the F2 and B2 conditions were blocked, it is possible that the observed differences stemmed from subjects anticipating different probes in the two conditions, and not from whether items were held as bound or separate features. F2 probes aligned with the memory array in only a single feature dimension, while the other dimension was held constant at a fixed value (Black/ Center of the screen). In this case, subjects could accomplish the task by generating a template encompassing all feasible matching probes. For instance, if the memory array contained a red item positioned in the upper-right quadrant and a blue item in the lower left, participants could devise templates of a red item situated at the center, a blue item at the center, a black item located in the upper-right quadrant, and a black item in the lower-left quadrant, all of which would require an ‘Match’ response in the F2 condition. In the B2 condition, in contrast, it was enough to compare the probe directly with the retained memory array without the need to recreate potential matching templates. If this were the case, bound items were retained in both conditions, and the variations in ERP between the F2 and B2 conditions reflected the process of template generation and maintenance.

Experiment 2 was designed to resolve the concern by replicating the results of Experiment 1 while presenting unpredictable probes that are identical between B2 and F2 conditions.

## 3 Experiment 2

The above study revealed significant differences in EEG signals recorded in central-parietal and frontal channels. As identical memory arrays were presented for both conditions, these dissimilarities suggested that the representation formats of objects within working memory were task-dependent. However, the anticipation of the different probes in the two conditions could contribute to varied cognitive processing during the working memory delay phase unrelated to the maintenance format. Experiment 2 aims to provide a conceptual replication with a similar paradigm, except that the probes were similar in both conditions, consisting of two items of distinct colors in different locations. Thus, the only difference between these two conditions was the judgment rule. In the B2 condition, participants were required to respond "Match" solely when one of the items in the probe matched one of the items in the memory array on both color and location (that is, conjunctions mattered). Conversely, in the F2 condition, participants were required to respond "Match" when the color or location of either item had been presented on the memory array.

### 3.1 Method

#### 3.2.1 Participants

Twenty-four healthy volunteers from the Hebrew University of Jerusalem participated in the study. Subjects were paid (∼$12/h) or given course credits for participation. All subjects had reportedly normal or corrected-to-normal sight and no psychiatric or neurological history. One subject was excluded due to noisy recordings. The remaining twenty-three subjects (Mean = 25.4 years old, SD = 3.2) consisted of 11 males and 12 females. The experiment was approved by the ethics committee of the Hebrew University of Jerusalem, and informed consents were obtained after the experimental procedures were explained to the subjects.

#### 3.2.2 Stimuli and Apparatus

Subjects sat in a dimly lit room. The stimuli were presented with Psychotoolbox-3 (http://psychtoolbox.org/) implemented in Matlab 2018 on a 57× 34.7 cm BENQ XL2411P (1024×768) monitor with a 144-Hz refresh rate. They appeared on a grey background at the center of the computer screen located 60 cm away from the subjects’ eyes.

Subjects performed a delayed (Yes/No) recognition test in 2 conditions (Figure 3a): Feature (F2), and Binding (B2). The memory arrays were identical to the items presented in Experiment 1. However, in Experiment 2, the probe arrays were the same in B2 and F2 conditions and each probe array contained two items (Figure 3a). One of the items matched the memory array in neither color nor location; the other item determined the probe type, which could be one of five types described in Table 2.

**Figure 3.**
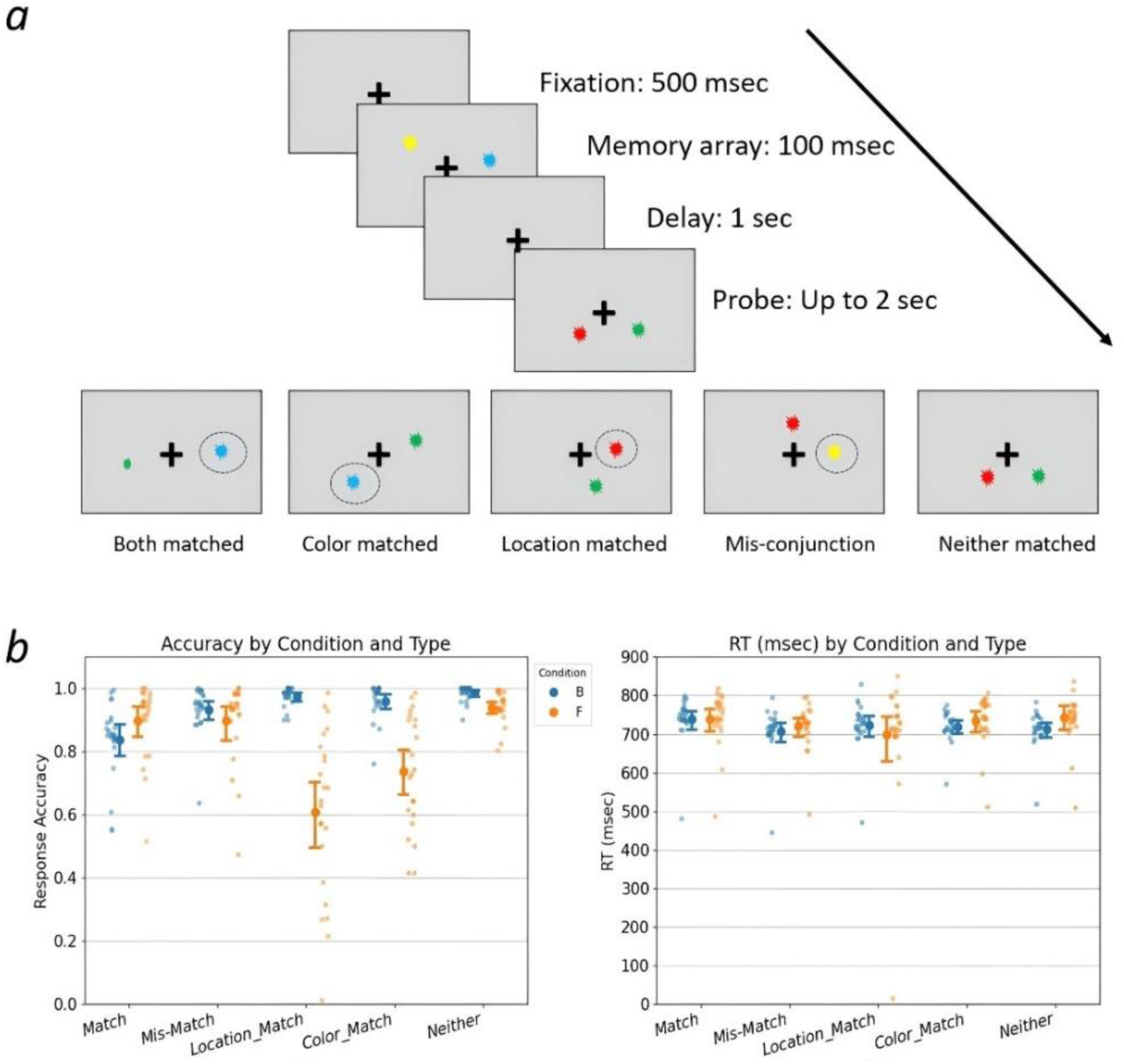
(a) Experiment 2 design. Top: Time course of one trial culminating in a "Neither" matched probe. Bottom: Illustration of probes from the five types. The probe is labeled relative to the array shown in the top panel. The circle around one of the stimuli is presented here to emphasize the item that defines the type of the trial but was not shown to the subjects. (b) Proportion of correct responses and reaction time for each type of probe in F2 and B2 conditions. Small dots represent individual subject means; large dots and error bars depict across-subject means and 95% confidence intervals.

**Table 2:** Description of probes used in Experiment 2.

| Probe type | Description | Correct Response in F2 condition (Trial number) | Correct Response in B2 condition (Trial number) |
| --- | --- | --- | --- |
| Matched | A probe with the same color and location as one of the items in the memory array. | Matched (70) | Matched (375) |
| Color-matched | A probe with the same color as one of the items in the memory array but in a new location. | Matched (70) | Non-matched (70) |
| Location-matched | A probe with the same location as one of the items in the memory array but with a new color. | Matched (70) | Non-matched (70 ) |
| Mis-conjunction | A probe with color and location both presented in the memory array but not in the same item | Matched (165 ) | Non-matched (165) |
| Neither-matched | A probe with color and location not presented in the memory array. | Non-matched (375) | Non-matched (70) |

#### 3.2.3 Experimental procedure

The procedure was similar to that of Experiment 1, with the following exceptions.

1. To allow for more trials in the experiment, a shorter fixation duration of 500 msec was used before the presentation of the memory array.
2. As the effect we found was shorter than 1 sec, only 1 sec was applied as delay duration.
3. The experiment encompassed only two conditions, each comprising a total of 750 trials spread across 5 blocks. The trial number for each probe type is shown in Table 2. To initiate each condition, a block of 120 practice trials was conducted, followed by five consecutive testing blocks. Each block took about 8 mins. The order of the two conditions was counterbalanced between subjects.

The only difference between the F2 and B2 conditions was the task demands. In the B2 condition, participants were required to respond "Matched" if one of the items matched an item in the memory array on both color and location. Successful execution of this task required participants to not only retain the individual feature values but also to recall the associations between color and location. Conversely, within the F2 condition, participants were required to respond “Matched" whenever the color or the location of one of the probe items was the same as one of the features presented in the memory array. For this task, participants had to retain the feature values without necessarily retaining the color-location relationships. Moreover, ignoring the conjunction information might reduce interference and facilitate the performance in this condition. For example, the Color-matched probe illustrated in Figure 3a requires a “Matched” response in the F2 condition because the color blue is present in the memory array. In contrast, the same probe requires a “Non-matched” response in the B2 condition because the blue item is in the wrong place. As both memory arrays and probes were the same between B2 and F2 conditions and no template of the Matched probe could be anticipated, ERP differences between the F2 and B2 conditions, occurring before the onset of the probe, would reflect the a cleaner “Task effect”.

#### EEG recording and data analysis

Response accuracy was entered into a Probe type ×Condition 5 ×2 two-way repeated-measures ANOVA. EEG data recording, preprocessing and analysis were the same as in Experiment 1(See Supplementary Table S2 for the deleted channels for each subject in Experiment 2).

### 3.2 Results

#### 2.4.1 3.2.1 Behavioral results

A Probe type × Condition, 5 × 2 repeated-measures ANOVA of response accuracy revealed a significant interaction between Probe Types and Condition, *F(4,88)* = 33.85, *p* < .001, *η2* = 0.61 (Figure 3b). Follow-up simple effects of the Condition factor revealed a significantly lower response accuracy in the F2 condition than the B2 condition on the *Location-Match probe* and *Color-Match probe*, but not for other probe types (See Supplementary Table S3). A similar ANOVA on RTs found a significant interaction between Probe type and Condition, *F(4,88) = 3.02, p = .022, η2 =0.02.* However, the effect size was small and no significant differences between conditions were found with follow-up simple effect analysis (See Supplementary Table S4). This suggests that the differences between conditions in Location-Match and Color-Match probes were not due to time-accuracy trade-offs. One of the explanations for the difficulty in detecting matched color (associated with a non-matched location) or matched location (associated with a non-matched color) in the F2 condition was interference from the other feature.

#### 2.4.2 3.2.2 Linear-Mixed model results

The F2 condition elicited a more positive voltage compared to the B2 condition in Centro-parietal electrodes, replicating the result of Experiment 1 (See Figures 4a and 4b). The centro-parietal cluster was accompanied by more negative voltage in F2 than B2 condition in surrounding channels.

**Figure 4.**
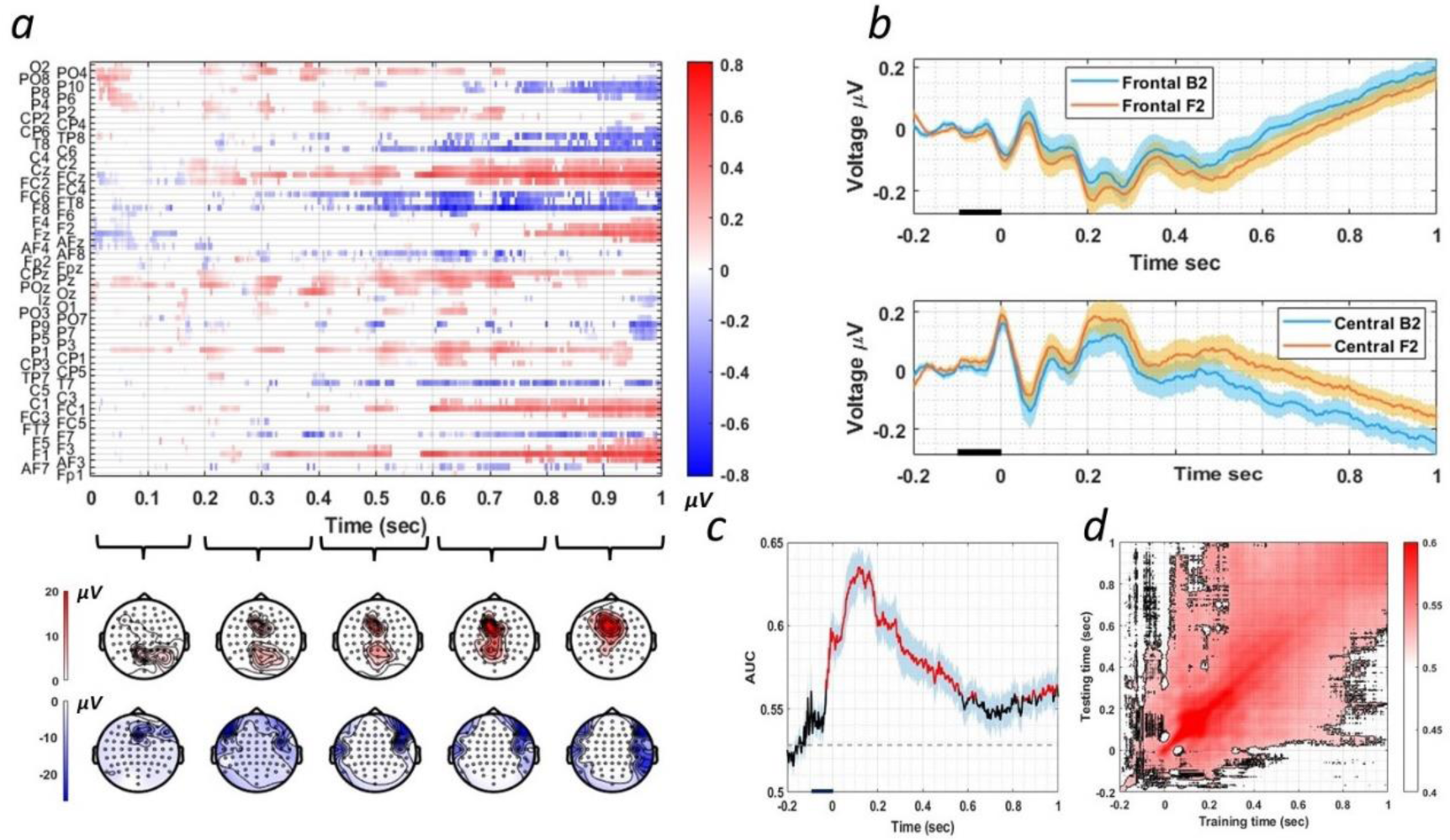
EEG results of Experiment 2. (a) Top Panel: Difference matrix between the F2 and B2 conditions, where significant (p < 0.05, FDR corrected) positive differences are displayed in red and negative differences in blue. The y-axis labels correspond to each channel, with differences shown for each time point. Bottom Panel: Difference topographies between the F2 and B2 conditions, with colors representing the sum of significant voltage differences (F2 - B2) over each 200 ms interval (p < 0.05, FDR corrected). The top row shows the positive differences, and the bottom row shows the negative differences. (b) Average event-related potentials (ERPs) triggered by the F2 and B2 conditions, averaged across subjects. The top panel displays ERPs from channels showing significant positive differences, while the bottom panel shows ERPs from channels showing significant negative differences. (c) Accuracy (AUC) of decoding the task (B2 vs. F2) in Experiment 2. Red segments denote AUC significantly higher (p < 0.01, FDR corrected) than the mean AUC during -100 to 0 ms (represented by the dashed horizontal line) relative to memory array onset. Shaded areas represent the standard error across subjects. (d)Temporal generalization matrix for decoding between B2 and F2 conditions, with testing time on the vertical axis and training time on the horizontal axis. The contour represents the significant cluster relative to 0.5, the chance level (p < 0.01, FDR corrected). Note that the AUC line in the (c) is the diagonal of the matrix in the (d).

#### 2.4.3 3.2.3 MVPA results

Significant time-resolved decoding between B2 and F2 conditions suggests that the temporal distribution of neural activity is different between conditions despite similar visual stimuli and anticipated probe arrays (Figure 4c). The classifier performance was found to be significantly above the chance level (AUC = 0.5) starting even before the onset of the memory array, peaking at 114 milliseconds and lasting through the whole delay period (*p < 0.01* FDR corrected). This early onset of decoding is not improbable, given that B2 and F2 conditions are presented in blocks, allowing subjects to prepare which task to perform before the onset of the memory array. To isolate the Task effect beyond preparatory effects, we compared the decoding accuracy to the mean AUC from -100 to 0 milliseconds before the onset of the memory array. Decoding was significantly higher than this baseline from 30 milliseconds before the onset of the delay, lasting until 550 milliseconds (*p < .01*, FDR corrected) and reemerged later during the delay period. These results suggest that the topographical distinction between conditions begins early during the encoding of stimuli, beyond overall differences between conditions. Figure 4d shows the temporal generalization of these results, with significance (*p < .01*, FDR corrected) relative to chance.

In summary, the results of Experiment 2 confirmed those of Experiment 1 – the requirement of maintaining bound objects reflects in less positive potentials in central electrodes. Importantly, since the same types of probes were presented in the B2 and F2 conditions, Experiment 2 helped eliminate the possibility that differences in the probes could account for the condition effects in Experiment 1. Taken together, the results of Experiments 1 and 2 support the hypothesis that the maintenance format in VWM is flexible rather than fixed, as required by changing task demands. Next, we aimed to extend this finding to other cases, specifically when location was not used as a task-relevant feature.

## 4 Experiment 3

In both Experiment 1 and Experiment 2, we focused on feature binding between location and color, which may be a unique case. Evidence suggests that objects can be automatically encoded alongside their spatial location. For instance, some studies have demonstrated that the seemingly irrelevant location of a stimulus can be decoded directly from EEG data during the delay period of a Visual Working Memory (VWM) task (Elsley & Parmentier, 2015; Foster, Bsales, Jaffe, & Awh, 2017; Olson & Marshuetz, 2005). EEG studies have further illuminated that spatial attention can be directed toward items in VWM even when the cue is non-spatial and the location is not relevant to the task (Eimer & Kiss, 2010; Kuo, Rao, Lepsien, & Nobre, 2009). Some research even suggests that the binding of non-spatial features might be mediated by their connection to spatial locations, possibly due to the fact that many neurons that are selective to visual features also exhibit a degree of spatial selectivity (Schneegans & Bays, 2017; Treisman & Gelade, 1980; Pertzov & Husain, 2014). Given these considerations, we wondered whether the patterns observed in Experiments 1 and 2 can be extended to situations involving the binding of two features without the direct involvement of location memory. Therefore, in Experiment 3, we investigated Color-Orientation binding, using the same experimental paradigm as in Experiment 2.

### 4.1 Methods

#### 4.1.1 Participants

Twenty-eight healthy volunteers from the Hebrew University of Jerusalem participated in the study. Subjects were paid (40NIS/h, ∼$12) or given course credits for participation. All subjects had reportedly normal or corrected-to-normal sight and no psychiatric or neurological history. Six subjects were excluded before analysis. One was due to chance performance in the memory tasks, while the other five were due to excessive noise in the recording based on the a priori prescribed criteria. The remaining twenty-two subjects (*Mean = 23.9 SD =2.43* years) consisted of 4 males and 18 females. The experiment was approved by the ethics committee of the Hebrew University of Jerusalem, and informed consents were obtained after the experimental procedures were explained to the subjects.

#### 4.1.2 Stimuli and apparatus

Subjects sat in a dimly lit room. The stimuli were created and presented with Psychotoolbox-3 (http://psychtoolbox.org/) implemented in Matlab 2018 on a 57×34.7 cm BENQ XL2411P (1024×768) monitor with a 144-Hz refresh rate. They appeared on a black background at the center of the computer screen located 60 cm away from the subjects’ eyes.

As in experiments 1 and 2, subjects performed a delayed (Yes/No) recognition test (Figure 5a). In both conditions, the memory array consisted of 2 colored Gabor gratings (6 cycles, contrast 0.7; Figure 5a). Each item subtended a visual angle of 1.6°×0.9°. One grating was positioned on the right side, while the other was located on the left side of the midline. The distance between the center of each Gabor grating and the screen center was 1.3°. Each grating was assigned a color from a set of four distinct colors: red (RGB 255, 0, 0), green (0, 255, 0), magenta (255, 0, 255), and blue (0, 0, 255) and an orientation from four possible angles (3, 49, 95, or 141 degrees) relative to the horizontal line, without repetition (i.e. each item had a unique color and a unique orientation within a given array).

**Figure 5.**
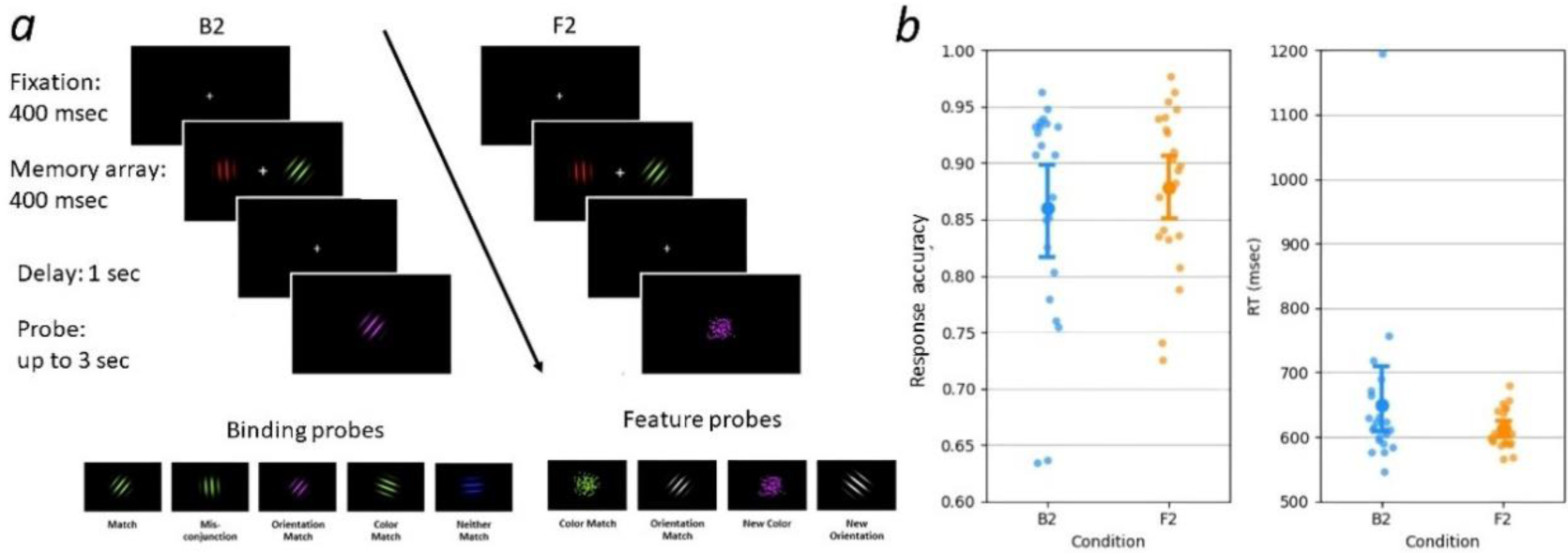
(a) Left panel: Example of a Binding trial with a “neither match” probe on the top, with examples of five Binding probes on the bottom. The Match state label of the probes in the bottom panel refers to the memory array shown in the top panel. Right panel: Example of a feature trial with a “New Color” probe, with examples of four Feature probes labeled relative to the memory array in the bottom. (b) Behavioral results of Experiment 3. The proportion of correct responses on the left panel and the reaction time on the right panel. Small dots represent individual subject means; large dots and error bars depict across subjects’ mean and 95% confidence interval.

#### 4.1.3 Experimental procedure

The procedure was the same as Experiment 2 with the following exceptions:

In Experiment 3 (Figure 5a), each trial started with a 400 msec fixation cross presented at the center of the screen. Next, the memory array appeared for 400 msec. This memory array presentation was 300 msec longer than that in Experiment 1 and Experiment 2 following pilot studies with a few subjects, indicating that a longer presentation period was necessary to effectively encode orientation and color compared to location and color.

Each condition began with 64 practice trials followed by 10 consecutive blocks of 72 trials of that condition, resulting in 720 trials in each condition. The trial numbers for each probe type are shown in Table 3. An equivalent number of matched and unmatched trials were incorporated within both conditions. Each block took about 5 minutes, and subjects were instructed to take a break between blocks. Odd subjects started with the Binding condition blocks followed by the Feature condition blocks, whereas even subjects performed the conditions in the reversed order.

**Table 3:** Description of probes used in Experiment 3.

| Probe type | Match State | Probe Description | Correct Response | Trial number |
| --- | --- | --- | --- | --- |
| Binding probe<br>(B2 condition) | Matched | Same grating as one of the items of the memory array in both color and orientation. | Matched | 360 |
|  | Color-matched | Grating with a color presented in the memory array, with an orientation did not appear in the memory array. | Non-matched | 60 |
|  | Orientation-matched | Grating with orientation present in the memory array, in a color that did not appear in the memory array. | Non-matched | 60 |
|  | Mis-conjunction | Grating with the color of one of the items in the memory array, combined with the orientation of the other item in the memory array. | Non-matched | 180 |
|  | Neither-matched | Grating matched neither the color nor the orientation of the items in the memory array. | Non-matched | 60 |
| Feature Probe<br>(F2 condition) | Color-matched probe | Cloud of dots, in one of the colors that appeared in the memory array. | Matched | 180 |
|  | New-color | Cloud of dots in a color absent from the memory array. | Non-matched | 180 |
|  | Orientation-matched | A grey Gabor grating with one of the orientations present in the memory array. | Matched | 180 |
|  | New-orientation probes | A grey Gabor grating with an orientation not present in the memory array. | Non-matched | 180 |

In the Binding condition (B2), a probe could take on one of five different types: Matched, Color-matched, Orientation-matched, Mis-Conjunction, and Neither- matched. These probes were presented at the center of the screen, sharing the same size and visual characteristics (cycle and contrast) as one of the Gabor gratings presented in the memory array (Table 3).

In the Feature condition (F2), there were four types of probes: Color-matched, New-color, Orientation-matched, and New-orientation. Color-matched and New-color probes were presented as a cloud of dots randomly located within a circle at the center of the screen with a diameter of 1.6°. Orientation-matched and New-orientation probes were grey Gabor grating, sharing the same size and visual characteristics (cycle and contrast) as the items presented in the memory array (Table 3).

#### 4.1.4 EEG and eye-tracker recording and data analysis

EEG data were recorded using the same procedure as in Experiments 1 and 2, while eye-tracking data were recorded with a desktop-mounted Eyelink 1000/2K infrared video-oculography system (SR Research Ltd., Ontario, Canada) at a sampling rate of 1000 Hz. A 9-point calibration procedure, followed by a validation stage (with acceptance criteria of worst point error < 1.5 visual degrees and an average error < 1.0 visual degrees), was applied before each block. A drift correction was then applied before each block started. Offline EEG and eye-movement synchronization were achieved by utilizing triggers sent from the stimulation computer via a parallel port. These triggers were simultaneously recorded by both the EEG recording system and the eye tracker.

Data preprocessing and analysis were done with the FieldTrip toolbox (version 20191213 http://www.fieldtriptoolbox.org/) implemented in Matlab 2018 (Mathworks, Natick, MA, USA). Preprocessing was applied to continuous data was similar to that used in experiments 1 and 2 except that subjects with over 4 channels deleted was excluded (See Supplementary Table S5 for the deleted channels for each subject) and an additional ICA procedure used to remove eye movement artifacts using data recorded by the eye tracker (Detailed in S4).

### 4.2 Results

#### 4.2.1 Behavior results

As shown in Figure 5b, accuracy and RT were comparable between the two conditions. Paired sample t-test revealed non-significant differences between Binding and Feature conditions both in accuracy (*Mdiff = -0.02, SE = 0.014, t(21)= -1.30, p = 0.21*), and in RTs (*Mdiff=38.02, SE=30.13, t(21)=1.26, p = 0.22)*.

#### 4.2.2 Linear-Mixed model results

The F2 condition elicited a more positive voltage compared to the B2 condition in Centro-parietal electrodes, replicating the results of Experiments 1 and 2. Additionally, similar to Experiment 2, more negative voltages were found in F2 than in B2 condition in surrounding channels (Figure 6a,6b).

**Figure 6.**
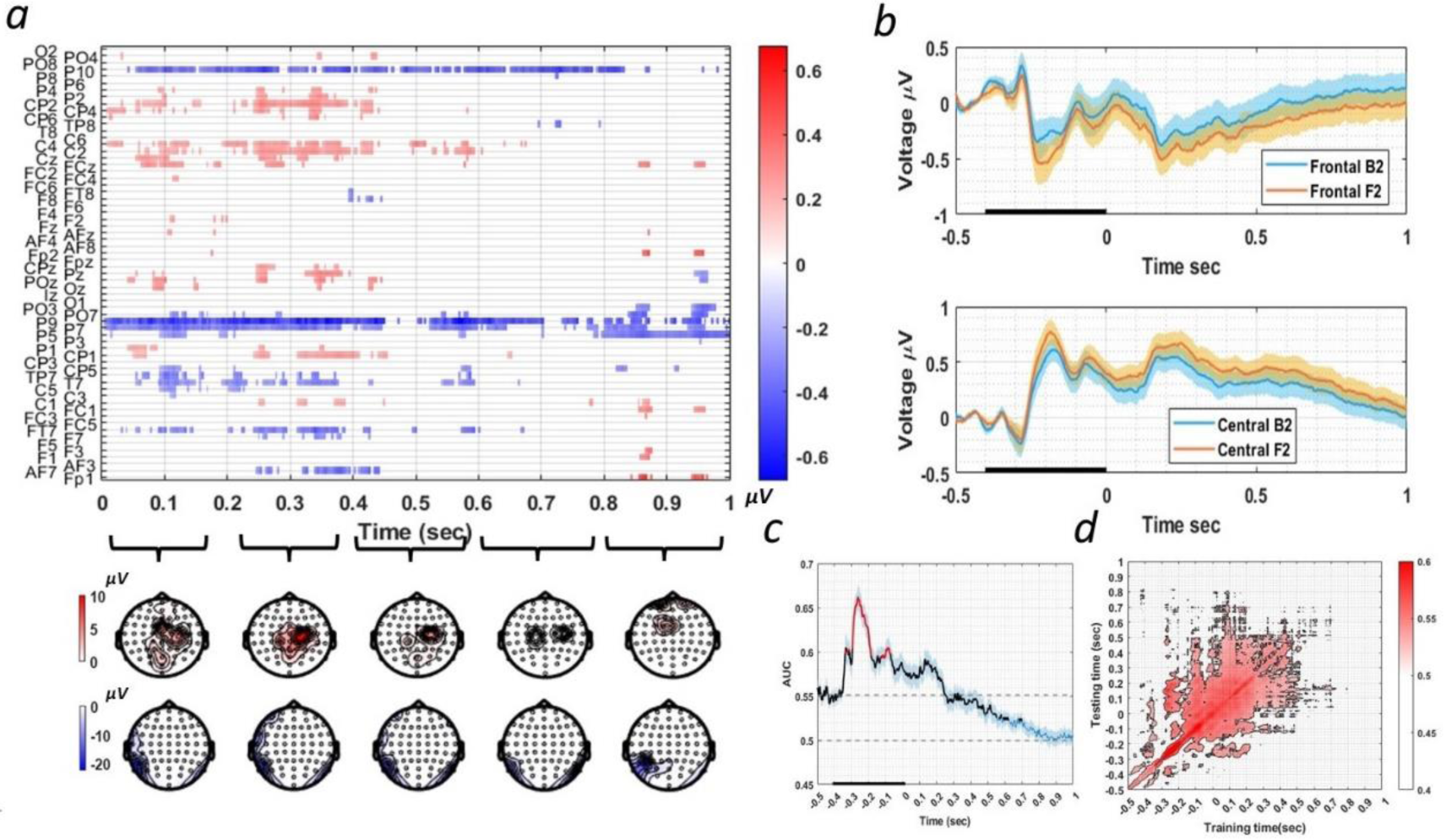
EEG results of Experiment 3. (a) Top Panel: Difference between the F2 and B2 conditions, where significant (p < 0.05, FDR corrected) positive differences are displayed in red and negative differences in blue. The y-axis labels correspond to each channel, with differences shown for each time point. As in other panels, Time 0 is the beginning of the delay period (note that the stimulus duration was longer than in Experiments 1 and 2). Bottom Panel: Difference topographies between the F2 and B2 conditions, with colors representing the sum of significant voltage differences (F2-B2) over each 200 ms interval. The top row shows the positive differences and the bottom row shows the negative differences (b) Event-related potentials (ERPs) triggered by the F2 and B2 conditions, averaged across subjects. The top panel displays ERPs averaged across channels with positive differences, while the bottom panel shows ERPs from channels with negative differences. (c) Accuracy (AUC) of decoding the task (B2 vs F2). Red segments denote AUC significantly higher (p < 0.01, FDR corrected) than the mean AUC during -100 to 0 (represented by the dashed horizontal line), relative to memory array onset. Black color denotes significant (p < 0.01, FDR corrected) AUC compared to 0.5 (represented by the bottom horizontal dash line). Shaded areas represent the standard error across subjects. (d) Temporal generalization matrix for decoding between B2 and F2 conditions with testing time on the vertical axis and training time on the horizontal axis. The contour represents significant accuracies relative to 0.5, the chance level (p < 0.01, FDR corrected). Note that the AUC line in panel (c) is the diagonal of the matrix in panel (d).

#### 4.2.3 MVPA results

AUC of decoding between B2 and F2 was found to be significantly above the chance level (0.5) starting100 msec before the onset of the memory array, peaking at 138 msec after the onset of the memory array (262 msec before the delay onset) and lasting until around 700 msec into the delay period (*p < 0.01* FDR corrected; Figure 6c). Temporal generalization analysis (Figure 6d) showed that classifiers trained on time points prior to stimuli onset could not decode the later maintenance period above chance. Meanwhile, the later component starting at the end of the memory array generalized for around 500 msec.

As in the previous experiments, we further compared the AUC to the mean of 0-100 sec prior to the memory array onset. A significant increase in decoding accuracy was found from 60 milliseconds following the onset of the memory array (340 msec before the delay onset) lasting until the end of array presentation (*p* < 0.01 FDR-corrected).

In summary, in Experiment 3, where location was not a task-relevant feature, we observed increased positivity in the F2 condition compared to the B2 condition at central channels. This is similar to the Task effect found in Experiments 1 and 2, where location was a task-relevant feature. This cluster emerged at the onset of the memory array and had a relatively short duration compared to the Task effect observed in the previous two experiments.

## 5 Discussion

In three experiments, we found significant differences in EEG signals between tasks requiring short term maintenance of visual features and tasks involving the maintenance of bound objects, when the visual input to be remembered (memory array) was identical across conditions. This provides evidence of task-dependent working memory processes and argues against the notion, implicitly embedded in previous literature, that a fixed default mode of item representation, either bound or not, is maintained in visual working memory.

In Experiment 1, we observed an effect of task-relevance of conjunction between location and color, characterized by increased positivity in centro-parietal channels when conjunction memory was not required (or increased negativity when conjunction memory was required). In Experiment 1, whereas the memory arrays were identical, different probes were used in the two conditions. This left open the possibility that the observed divergent EEG signals could be attributed to the process of creating a template encompassed all possible matched probes in Feature condition, which was not possible in the Binding condition. Therefore, in Experiment 2, subjects were presented with identical memory arrays and identical probes across conditions. Nevertheless, a similar pattern, strongly correlated with the Task effect revealed in Experiment 1, was replicated, suggesting that the observed Task effect in Experiment 1 went beyond the process of predicting the probe. In Experiment 3, we extended the Task effect from location-color binding, which may be a unique form of binding, to binding between color and orientation, where location is not a task-relevant feature.

A previous visual search study (Berggren & Eimer, 2018) provided a hint that the brain represents information in a format compatible with the test demands. In that study, subjects answered whether the search display contained one of two target items held in memory. When the search array included only one object with a matching feature, targets and incorrect conjunction objects elicited identical N2pc components and sustained posterior contralateral negativity (SPCN). The N2pc is assumed to index a shift of spatial attention to the location of the potential target (Eimer, 1996; Kiss, Van & Eimer, 2008), and the SPCN is assumed to index attentional activation of VWM representations of the potential target (Jolicoeur, Brisson, & Robitaille, 2008). These results suggest that in this condition only features were used as searching templates. In another condition, it was insufficient to detect a certain feature, as all objects had target-matching features, and both a target and an incorrect conjunction object could be present in the same display. In this case, the target evoked a larger N2pc than the incorrect conjunction objects, and only targets elicited SPCN components. Therefore, in this condition, a bound object, rather than an isolated feature template, was used by the subjects. Taken together, the results of this study are compatible with the possibility that VWM templates guiding attention in visual search were encoded or used flexibly, to effectively distinguish match and no-match probes, based on the task demand. However, this study specifically examined the impact of the search template provided at the beginning of the experimental condition on the response to the subsequent probe. Though the results might reflect differences in mode of maintenance of the search template, no direct measurements were taken during the encoding and maintenance periods in VWM.

In another study, Woodman and Vogel (2008) more directly addressed task-specific encoding and retention of visual stimuli during a delay period. In their study, a lower contralateral delay activity (CDA), considered to be an EEG marker of memory load, was observed when only color was task-relevant compared to the case where both color and orientation were relevant. By providing evidence that features within a bound item can be selectively retained or suppressed based on task requirements, these results challenged the notion that VWM representations are exclusively stored as integrated objects. Moreover, this flexibility is not only demonstrated when subjects have prospective knowledge of the task-relevant feature but is also observed when task relevance of stimuli was informed after the presentation of the memory array by retro cues (Niklaus, Nobre, & Van Ede, 2017 ; Park, Sy, Hong, & Tong, 2017). However, in the feature condition in those paradigms, only one feature had to be maintained, enabling subjects to ignore the other feature. In contrast to these studies, which manipulated which features had to be maintained, in our paradigm both features had to be maintained in both conditions, while the task-relevance of conjunction between features was directly manipulated. Our findings unveil an additional layer of flexibility, indicating that the maintenance of conjunction information itself, presumably the format of representation, as the bound object or separate features, can also be influenced by the task demands.

The finding that objects are retained either as separate features or as bound items based on task requirements may explain some of the seemingly conflicting results in the literature, although not all of them. This is based on the idea that although two tasks are similar in their explicit instructions, subtle differences in the relationship between the memory array and the probe might affect the way that information is maintained in memory. For example, Luck and Vogel (1997; extended in Vogel, Woodman, & Luck, 2001) observed that the ability to detect a change in a stimulus array of a given set size is the same regardless of the number of features. In contrast, Wheeler and Treisman (2002) found memory capacity is determined merely by the number of total features when two features from the same dimension are combined into one object. In Luck and Vogel’s study, each item presented in the memory array contained features randomly selected from all possible feature values with replacement (e.g., more than one item could be red) and the non-match probe could include an erroneous conjunction of features that appeared separately in the memory array. In contrast, in Wheeler and Treisman’s study (Experiments 1 and 2), items on the memory array were generated by selecting from possible feature values without replacement (e.g. only one item could be red). Moreover, to generate a non-matched probe, the probe included a feature that had not been used by any item in the memory array. This design, where no feature-value repetition was allowed in a single memory array, can be also found in other studies arguing that delayed change detection performance was hindered by increasing the number of features to be remembered for each object (Oberauer & Eichenberger, 2013). We suggest that in designs like Luck and Vogel’s, bound objects have to be maintained to detect a non-match probe, while in designs like Wheeler and Triesman’s, remembering a list of all features in the memory array is sufficient to distinguish between match and non-match probe. This may induce different “strategies” for maintaining information. The current findings, therefore, draw attention to subtle details between experimental designs which could have an impact on the information that is maintained in memory.

In all three experiments, the Feature condition consistently produced a higher central positivity during the Feature condition compared to the Binding condition. In Experiment 1, we also found a stronger central positivity in the more demanding B4 condition, compared to the B2 condition (the Load contrast), which was also shown in other studies (Mecklinger & Pfeifer, 1996; Ruchkin & Johnson, 1990). The higher positivity in the feature conditions (F2) compared to the binding conditions (B2) could thus indicate the maintenance of a larger number of units (i.e., 4 feature values vs. 2 bound objects) during the delay period of the Feature condition compared to the Binding condition. However, the topographical distribution revealing the Binding effect (B4 vs. B2) and the Load effect (F2-B2) in Experiment 1 are not entirely the same, and no above-chance cross-decoding between the Task effect and the Load effect was found. Therefore, the Task effect might reflect some additional processes unrelated to the increased number of units maintained in working memory during the delay.

In Experiment 2, performance in the Feature conditions dropped significantly compared to the Binding condition. This decline was mainly due to failures in detecting matched items when only one feature matched, possibly indicating some preservation of conjunction information. The identical probes in both conditions (as opposed to the distinct probes in Experiment 1) might have reduced the manipulation’s effectiveness. Alternatively, the relative worse performance of the Feature condition could be due to the interference of task-irrelevant feature dimension. That is, subjects may have stored the features separately, but the response conflict (match on one dimension and non-match on the other dimension) hindered performance This explanation is supported by the strong Task effect in ERPs observed in Experiment 2. Another possibility is that the poorer performance and increased positivity in central channels reflect a higher working memory load in the Feature condition. Further studies are needed to determine the correct explanation.

Another limitation of the current study is that the task effect was identified through contrasting or decoding between blocks. The block design allowed for a more sensitive detection of the effect as it did not require subjects to frequently switch tasks. We found some above-chance decoding of conditions even before the onset of the memory array when subjects viewed the fixation cross and prepared to encode the array. This might reflect tonic differences between blocks due to task requirements or other long-lasting factors. To exclude this possibility, future studies could employ a paradigm that informs the task relevance of conjunction on a trial-by-trial level with pre-cues. Moreover, the current results suggest that task effects on visual working memory (VWM) processes could be present as early as sensory encoding (Boettcher, Gresch, Nobre, & Van Ede, 2021). Whether such manipulation can be implemented after the encoding stage could be explored using paradigms with retro-cues, where the task is informed to subjects after the memory array presentation.

To conclude, our study extends the view that VWM should not be construed as a fixed storage of sensory input, but a dynamic system actively engaging in the task and guiding behaviors (Boettcher, Gresch, Nobre, & Van Ede, 2021; Lee, Kravitz, & Baker, 2013). It provides a novel piece of evidence that not only which features are maintained but also the format of objects represented in the working memory, either as bound objects that integrate different feature dimensions or as separated features, is task-dependent. Moreover, the emerging flexibility of working memory and its sensitivity to subtle changes in the task design provide possible explanations for the apparent conflict between previous results regarding this question.

## 8 Supplementary

**Table S1:** Interpolated channels for each subject in Experiment 1.

| **Subject** | **Interpolated Channels** |
| --- | --- |
| S01 | FC5, F6 |
| S02 | AFz |
| S04 | AFz, FC5 |
| S05 | AF8, Iz |
| S06 | AFz |
| S07 | AF8, Iz |
| S08 | PO3, CP1 |
| S09 | AF8, Iz |
| S12 | CP3, P1 |
| S14 | AF7, F7, P9 |
| S16 | POz, Fp2 |
| S17 | T7, TP8, P9 |
| S19 | F8 |
| S20 | CP3, T7 |
| S23 | T7, CP3 |

**Table S2:**
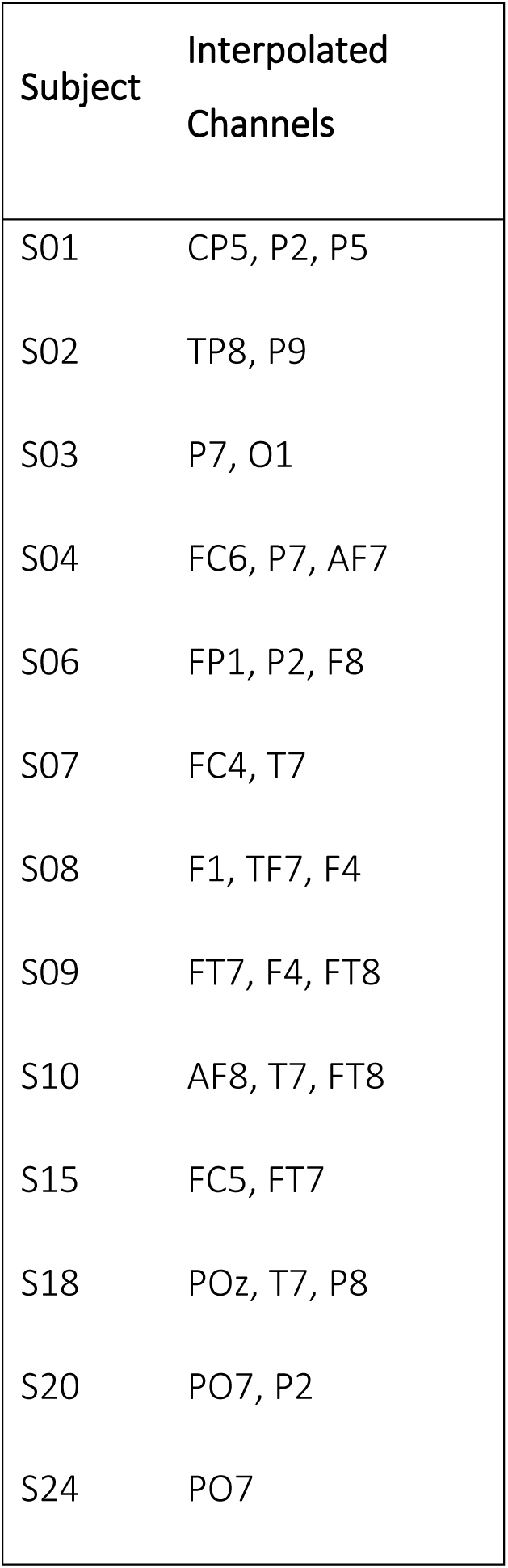
Interpolated channels for each subject in Experiment 2.

| Subject | Interpolated Channels |
| --- | --- |
| S01 | CP5, P2, P5 |
| S02 | TP8, P9 |
| S03 | P7, O1 |
| S04 | FC6, P7, AF7 |
| S06 | FP1, P2, F8 |
| S07 | FC4, T7 |
| S08 | F1, TF7, F4 |
| S09 | FT7, F4, FT8 |
| S10 | AF8, T7, FT8 |
| S15 | FC5, FT7 |
| S18 | POz, T7, P8 |
| S20 | PO7, P2 |
| S24 | PO7 |

**Table S3:**
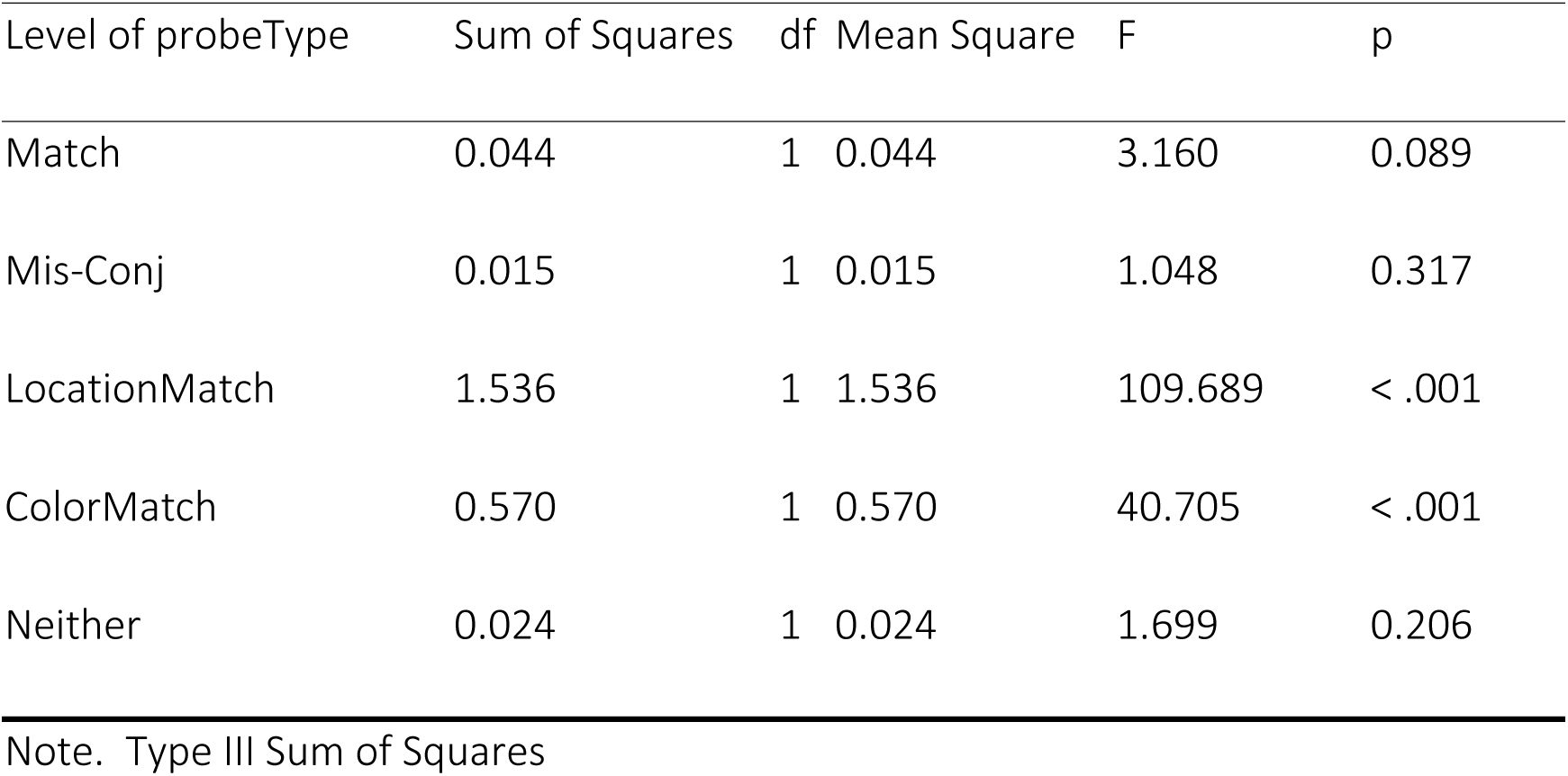
Simple Main Effects on Response Accuracy - Condition.

**Table S4:** Simple Main Effects on RT-Condition.

| Level of probeType | Sum of Squares | df | Mean Square | F | p |
| --- | --- | --- | --- | --- | --- |
| Match | 2.867 | 1 | 2.867 | 1.346e-4 | 0.991 |
| Mis-Conj | 1679.753 | 1 | 1679.753 | 0.079 | 0.782 |
| LocationMatch | 7397.995 | 1 | 7397.995 | 0.347 | 0.562 |
| ColorMatch | 1289.365 | 1 | 1289.365 | 0.061 | 0.808 |
| Neither | 10319.368 | 1 | 10319.368 | 0.484 | 0.494 |

**Table S5:** Interpolated channels for each subject in Experiment 3.

| Subject | Channels |
| --- | --- |
| S02 | Pz, FC4, P2, T7 |
| S03 | AF7, FC5 |
| S06 | FC6, P6 |
| S07 | AF7, FC5 |
| S08 | TP8, T7, AF8, AF7 |
| S09 | C4, CP6 |
| S10 | Oz, CP5, POz |
| S17 | C4, CP6 |
| S21 | FT7, T7, FT8, AFz |
| S22 | P3, Cz, AF8 |

## S4: Artifact Removal during Pre-processing Guided by EOG or Eye tracker for Experiment 3

During preprocessing, as in experiments 1 and 2, EEG and EOG signals were first filtered with Butterworth zero-phase (forward and reverse filter) bandpass filter of 0.1– 180 Hz and then referenced to the nose channel. Extremely noisy or silent channels, which contributed more than 20% of all artifacts (Criteria: more than 100μV absolute difference between samples within segments of 100 msec; absolute amplitude > 100μV) were deleted. No more than 2 adjacent channels or 4 channels in a single subject were deleted. Next, data were re-referenced to an average of all remaining EEG electrodes.

As a longer duration of the memory array was applied in Experiment 3 compared with previous two experiments, more eye-movements especially, saccades in the horizontal direction were found.

To find Independent Component Analysis (ICA) components that accurately capture eye-movement artifacts, we first combined segments from both the Binding (B) and Feature (F) conditions to create the training dataset. This dataset included peri-stimuli segments (from the onset of the memory array until the subject’s response), peri-saccade segments (with 30 samples before and 20 samples after saccade onset), and blink segments (encompassing 150 samples before blink onset and after blink offset). To improve the representation of eye-movement artifacts, we intentionally increased the proportion of saccades and blinks by duplicating these segments, resulting in up to 1000 segments in each condition(Keren et al., 2010). For the stimuli segments, we used every other segment to reduce computation. Therefore, we subtracted the mean of each segment from every data point. and high-pass filtered with a filter (4th-degree non-causal Butterworth filter with a Hamming window) with a cutoff frequency of 1.5 Hz. This way of training ICA was shown to improve the removal of eye movement related artifacts (Dimigen, 2020).

For fourteen subjects with eye-tracker data, the detection of blink and saccade onsets for the construction of the training data relied on the eye-tracker recordings. To identify saccades, we employed a previously published algorithm designed to detect and quantify binocular saccades within the unsegmented eye tracker data (Engbert and Kliegl, 2003). Saccade onsets were defined as time points at which the absolute eye velocity exceeded 6 standard deviations from the mean. Subsequently, we segmented the data into 80-millisecond epochs, starting 30 milliseconds before the onset of each saccade.

For eight subjects without available eye-tracker data, we identified saccades and blinks from the EOG channels offline. A bipolar vertical EOG (VEOG) channel was calculated as the difference between VEOGS and VEOGI. Similarly, a bipolar horizontal EOG channel (HEOG) was calculated as the difference between the left (HEOGL) and right (HEOGR) horizontal EOG differences. Following previous recommendations (Croft and Barry, 2000; Elbert et al., 1985;Keren, Yuval-Greenberg, & Deouell, 2010, we also derived a “radial” electro-oculogram channel (REOG) for the analysis of spike potentials (SPs) , defined as the average of all EOG channels referenced to Pz:

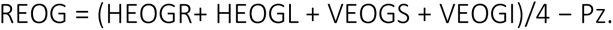

We then band-passed the REOG signal to 30-100 Hz using a 6th-degree non-causal Butterworth filter. SPs, indicating the onset of saccades, were identified as time points exceeding 2.6 standard deviations from the mean of this REOG signal. This threshold was determined as it yielded nearly identical segments as those identified by the eye tracker (in cases where both EEG and eye tracker data was available). To identify blinks, we applied a Butterworth zero-phase bandpass filter (forward and reverse filter) with a range of 1-10 Hz to the VEOG bipolar channel. Time points exceeding 2 standard deviations from the mean of this VEOG channel, along with a 100-millisecond interval before and after, were marked as blinks. Subsequently, a visual inspection of the data was conducted to ensure the precise identification of all saccades and blinks.

We manually identified Independent Component Analysis (ICA) components that represented muscle, blink, or eye-movement artifacts, based on their temporal profiles and scalp topography. These selected components were subsequently removed from the original preprocessed data (i.e. without the 1.5 Hz high pass filter used for the training data) .

Time points that exceeded 12 standard deviations from the mean for each condition of the corresponding channel were deleted, along with an additional 200 milliseconds before and after these points. This approach aimed to encompass the subthreshold beginnings and ends of artifact events. Following this automated procedure, a visual inspection of the data was conducted to identify any rare artifacts that might have been missed by the automatic method. Finally, previously removed channels were reconstructed by means of interpolation based on neighboring electrodes. The specific electrodes used for interpolation are presented in S3.

The data was next divided into segments of 1500 milliseconds each, starting 100 milliseconds before the onset of the memory array and ending at the appearance of the probe (1400 milliseconds after the memory array onset). The average of the 100 milliseconds preceding the onset of stimuli for each trial was calculated and subtracted from all data points within each segment. The data then underwent a Butterworth zero-phase lowpass filter, with a cutoff frequency of 20 Hz. Finally, the data was down- sampled to 256 Hz for further analysis.

